# Mapping the miRNA Landscape of Gallbladder Cancer: Genome-Wide Insights and Functional Implications

**DOI:** 10.1101/2025.09.29.679165

**Authors:** Sonika Kumari Sharma, Garima Singh, Simran Mathur, Deepanshu Aul, Puneet Gupta, Vinay Kumar Singh, Satyendra Kumar Tiwary, Samarendra Kumar Singh

## Abstract

Gallbladder cancer (GBC), the sixth most aggressive gastrointestinal cancer, exhibits complex molecular dysregulation involving microRNAs (miRNAs) that drive tumor progression. This study has comprehensively profiled genome-wide miRNA expression in GBC cells revealing 971 known miRNAs, and 27 novel miRNAs. We identified 388 significantly differentially expressed (DE) miRNAs (191 upregulated and 197 downregulated), providing valuable insights into the various complex molecular mechanisms underlying GBC pathogenesis. These DE miRNAs target key biological processes, including hormone-induced signaling pathways, cell adhesion molecules, apoptosis, ferroptosis, and cancer-associated pathways, particularly those linked to gastric cancer. Profiling of DE miRNAs in GBC cells presents opportunities for novel miRNA-based diagnostic and prognostic approaches in GBC. Notably, among the DE miRNAs, the miR-17∼92 cluster was significantly suppressed in GBC cells. We also show that miR-17∼92 cluster targets and destabilizes an S-phase licensing factor Cdt2 (an oncogene) in GBC cells. miR-17∼92 mediated negative regulation of Cdt2 induces S-phase arrest, leading to checked proliferation and invasive capabilities of GBC cells specifically, without any significant effect on non-cancerous cells, presenting miR-17∼92 as a potential miRNA therapeutic for GBC.

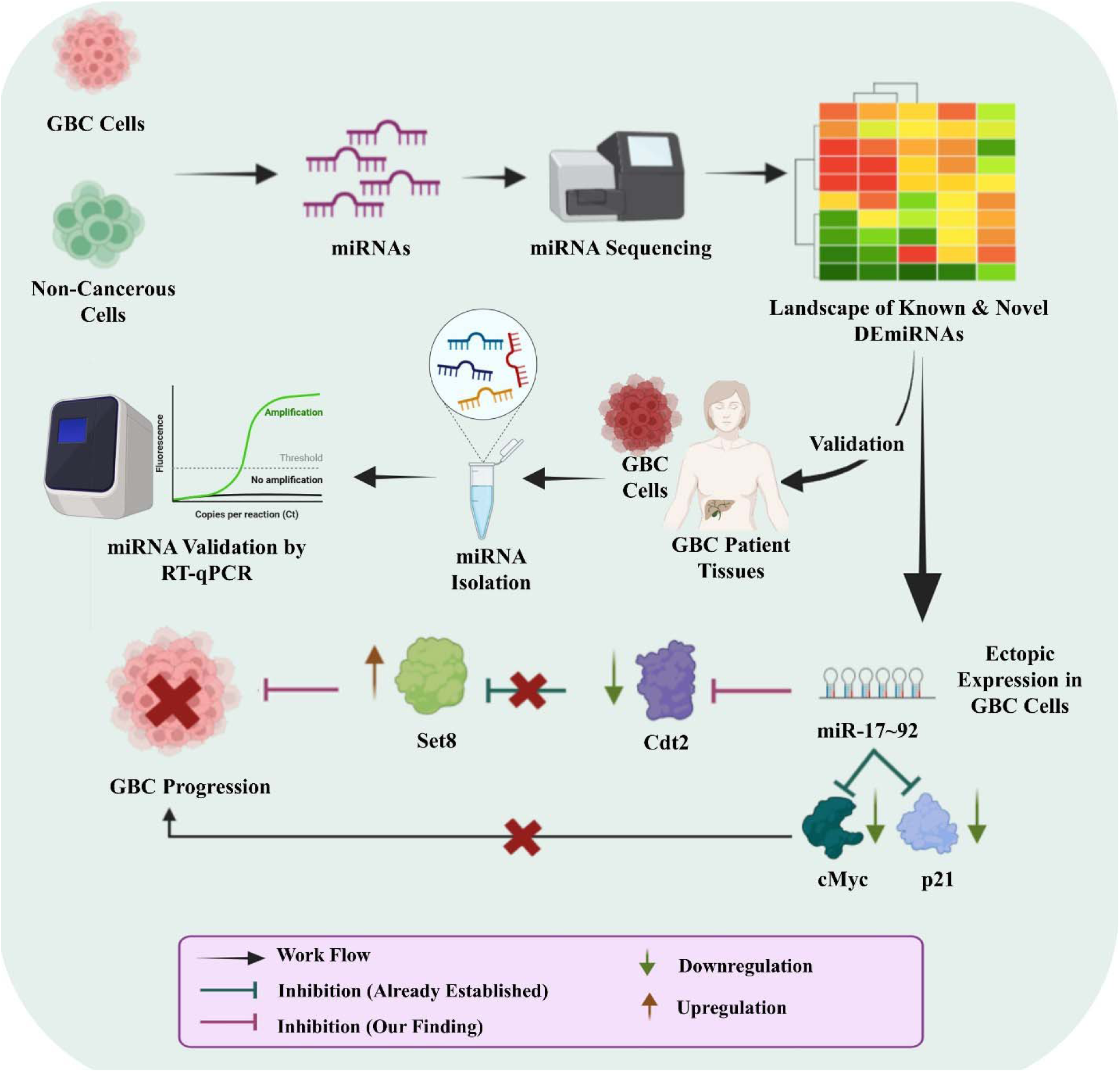
Graphical Abstract.

## Introduction

Gallbladder cancer (GBC) is the 6^th^ most common gastrointestinal cancer, accounting for 0.6% of global cancer burden (122,462 cases) and 0.9% (89,031 cases) of total cancer death (1). Being highly malignant with notable early-stage lymph node metastasis makes this disease quite aggressive. The cases have been predominantly reported in developing and underdeveloped countries of South-Central Asia, Northern Africa and South-America. Females and elderly people are more prone to GBC, than males and younger people respectively (1). The various etiological factors which could lead to higher rates of gallbladder cancer are gallstones, acute infection, genetically predisposed family history, gallbladder polyps, obesity etc. (2–4). It does not show any specific symptoms till cancer advances to critical stage where surgical intervention (cholecystectomy) is the only curative option available. The recurrence rate of the cancer is quite high even after cholecystectomy (5). There are various factors responsible for the oncogenesis of this cancer, like mutation in p53, p21, p27, Smad4, PTEN, FHIT genes etc. The above events lead to dysregulated cell cycle (6,7), which is the hallmark of oncogenic transformed cells. Therefore, exploring the cell cycle factors that are mis-regulated in GBC could open up new avenues for exploring therapies against this deadly cancer.

Cdt2 (Cdc10-dependent transcript 2)/DTL, an essential mammalian protein, is a master coordinator of cell cycle progression (for S-phase entry) and genomic stability. It is a substrate adaptor protein for CRL4 (Cullin RING Ligase Complex 4) E3 ubiquitin ligase system and targets critical cell cycle factors like Cdt1, p21 and Set8 for their ubiquitin mediated proteasomal degradation (8). Set8 (a monomethyltransferase) is a licensing factor which monomethylates H4K20 and initiates the replication event at the origin followed by loading of ORC, Cdc6, Cdt1 and MCM2-7 in G1 phase of the cell cycle. During S-phase, p21 destabilizes CDK2, which is an essential protein for the progression of replication initiation and S-phase progression. Hence, CRL4^Cdt2^ E3 ligase mediated degradation of Cdt1, Set8 and p21 is essential for S-phase entry (9). Cdt2 in turn gets degraded by CRL1^FBXO11^, CRL4^DDB2^ and APC/C-Cdh1 at the end of S-phase which is important for the stabilization of Cdt1, Set8 and p21 (10). In many cancers, such as lung, cervical, colon, hepatocarcinoma, etc. Cdt2 is reported to be overexpressed which leads to re-replication and hyperproliferation of the cancerous cells (11). Recently, we showed that microRNA miR-34a destabilizes Cdt2 which in turn suppresses proliferation and metastasis of cervical cancer cells (12).

MicroRNAs are 17-25 nucleotides long, small non-coding single stranded RNA molecules that epigenetically regulate ∼60% of the total genes at post-transcriptional level. It works by binding mostly to 3’UTR of target mRNA (13), leading to either degradation of target mRNA or translational inhibition, depending upon the degree of complementarity. By regulating over 30% of the protein coding genes, miRNAs are involved in regulation of many essential cellular processes such as differentiation, development, proliferation, apoptosis, etc. (14,15). In cancer and various diseases, dysregulation of miRNAs contributes to abnormal cellular processes. MiRNAs, therefore, can either be oncogenic or tumor suppressor, depending upon the pathway they affect. The role of miRNAs is minimally explored in gallbladder cancer progression, leaving a large knowledge gap in the area. To address this, we performed the first-ever genome-wide miRNA profiling in gallbladder cancer cell line. Our analysis revealed significant differential expression of 388 miRNAs (compared to a non-cancerous cell line control), providing novel insights into the molecular mechanisms underlying this malignancy. The miRNA profiling, also led to the discovery of novel miRNAs which were not reported earlier. Taking the study forward, we validated some of the differentially expressed miRNAs in both GBC cell lines as well as GBC patient tissue samples, which opens up new experimental avenues deconvoluting the dynamics of these miRNAs in GBC pathogenesis. Our results further elucidate a mechanistic insight of miR-17∼92 mediated gene regulation of an oncogenic cell cycle factor Cdt2 in GBC cells, suggesting the potential therapeutic implication of miR-17∼92 in GBC.

## Materials and methods

### Cell culture

The human gallbladder cancer cell lines, NOZ was a provided by Dr. Srikanth, (Regional Centre for Biotechnology, Faridabad, India) and G415 was a gift from Dr. Amit Dutt (ACTREC, Mumbai, India). The human embryonic kidney cell line, HEK293T was purchased from the National Centre for Cell Science (NCCS, Pune, India). The cells were cultured in Dulbecco’s Eagle Modified Medium (DMEM; Gibco #12800-017) supplemented with 10% Fetal Bovine Serum (FBS; Gibco #10270-106) and 1% Penicillin-Streptomycin (Pen-Strep; Gibco #15140-122) at 37°C, with 95% humidity and 5% CO_2_ (unless mentioned otherwise).

### RNA isolation and miRNA sequencing

NOZ and HEK293T cells were cultured till they reached 70-80% confluency. Total RNA was isolated using miRVana^TM^ microRNA isolation kit (Invitrogen #AM1560) and quantified using Nanodrop spectrophotometer. RNA integrity and quality were assessed using TapeStation 2700 system (Agilent Technologies), yielding RNA Integrity Number (RIN) of 9.3 for both cell lines. Small RNA sequencing libraries were prepared from 1 µg of quality control (QC) passed total RNA using Illumina NEBNext® Multiplex small RNA library prep kit for Illumina (NEB #E7560S) as per manufacturer’s instruction. The enriched libraries were then purified using Spin column followed by AMPureXP bead-based size selection. The purified and size selected libraries were checked with Qubit and real-time PCR for quantification that were further analyzed for size distribution on 4200 TapeStation system (Agilent Technologies) using high sensitivity D1000 Screen tape (Agilent #5067-5582). The Quantified libraries (18-40 bp) were pooled and were subjected to single-end 50 bp sequencing on Illumina platform with SE50 sequencer.

### miRNA sequencing data analysis

Small RNA sequencing generated flow-cell images were base called into sequenced reads in FASTQ (fq) files by Illumina CASAVA v1.8 and determined the Phred quality scores to assess base-calling accuracy and read quality. The sRNA reads were further filtered to eliminate low-quality reads, and analyzed for length distribution to characterize the reads. The High-quality reads were then aligned to the human genome (GRC38_p14) using Bowtie software to map the sRNA reads. The mapped reads were then genome-wide represented for their chromosomal distribution using Circos. Known miRNAs were analyzed using miRBase miRNA database and novel miRNAs were predicted using the miREvo and miRDeep2. The read counts were normalized using TPM (transcript per million) method to ensure comparability. Differential expression analysis of miRNAs was performed using DESeq2 and functional enrichment of miRNA targets was conducted using Cluster Profiler package and g-profiler. The miRNA-mRNA interactome was constructed using the software Cytoscape_v3.10.3.

### Sample Collection

The histopathology confirmed tissue samples were collected from the gallbladder cancer patients (GBC; experimental group) and control gallbladder tissue samples were collected from patients with gallstone (GS; control group) admitted at Department of General Surgery, Institute of Medical Sciences, Banaras Hindu University, Varanasi. The collected tissue samples were stored in RNAlater (Invitrogen #AM7020) at –80 till further processing.

### Plasmids and miRNAs

The pcDNA3.1 vector (used as an empty control) was purchased from Addgene (#128034) and miR-17∼92 cloned in pcDNA3.1 vector was kindly provided by Dr. Joshua T. Mendell (16). Flag-tagged Cdt2 (Flag-Cdt2) expressing plasmid was generously gifted by Dr. Anindya Dutta’s Lab (University of Virginia, USA) (17).

### Transfection

HEK293T, NOZ and G415 cells were seeded in a 6-well culture plate at a density of 0.8X10^5^ cells per well. Post 24 hours of incubation, cells were transfected with 4 µg of either control plasmid, miR-17∼92 or/and Flag-Cdt2 (2 µg) using TurboFect^TM^ (Thermo Fischer Scientific #R0534) according to manufacturer’s protocol. The cells were then incubated at 37°C, with 95% humidity and 5% CO_2_ for 24-48 hours before they were harvested for further experiments.

### Real-Time Reverse Transcription PCR (qRT-PCR) For Mature miRNAs Quantification

Patients tissue samples (75-100 mg) were homogenized in TriZol (Thermo Scientific #15596026) and total RNA was extracted from tissue samples as wells as from cell lines. To quantify the mature miRNA expression, two step RT-qPCR was performed. Reverse transcription (RT) of mature miRNAs (miR-100-5p, miR-335-3p, miR-10b-3p, miR-10b-3p and miR-218-3p) were carried out with sequence specific stem-loop RT primers using the 1^st^ strand cDNA synthesis kit (Takara Bio #6110A) according to manufacturer’s instructions. Next, qPCR was performed to quantify the expression of mature miRNA (cDNAs prepared in previous step) using TB Green Premix Ex TaqTM II kit (Takara Bio #RR820A) according to manufacturer’s instructions using the cDNA specific forward primers and universal reverse primer (Table 1) with U6 snRNA serving as the endogenous control.

**Table 1:**
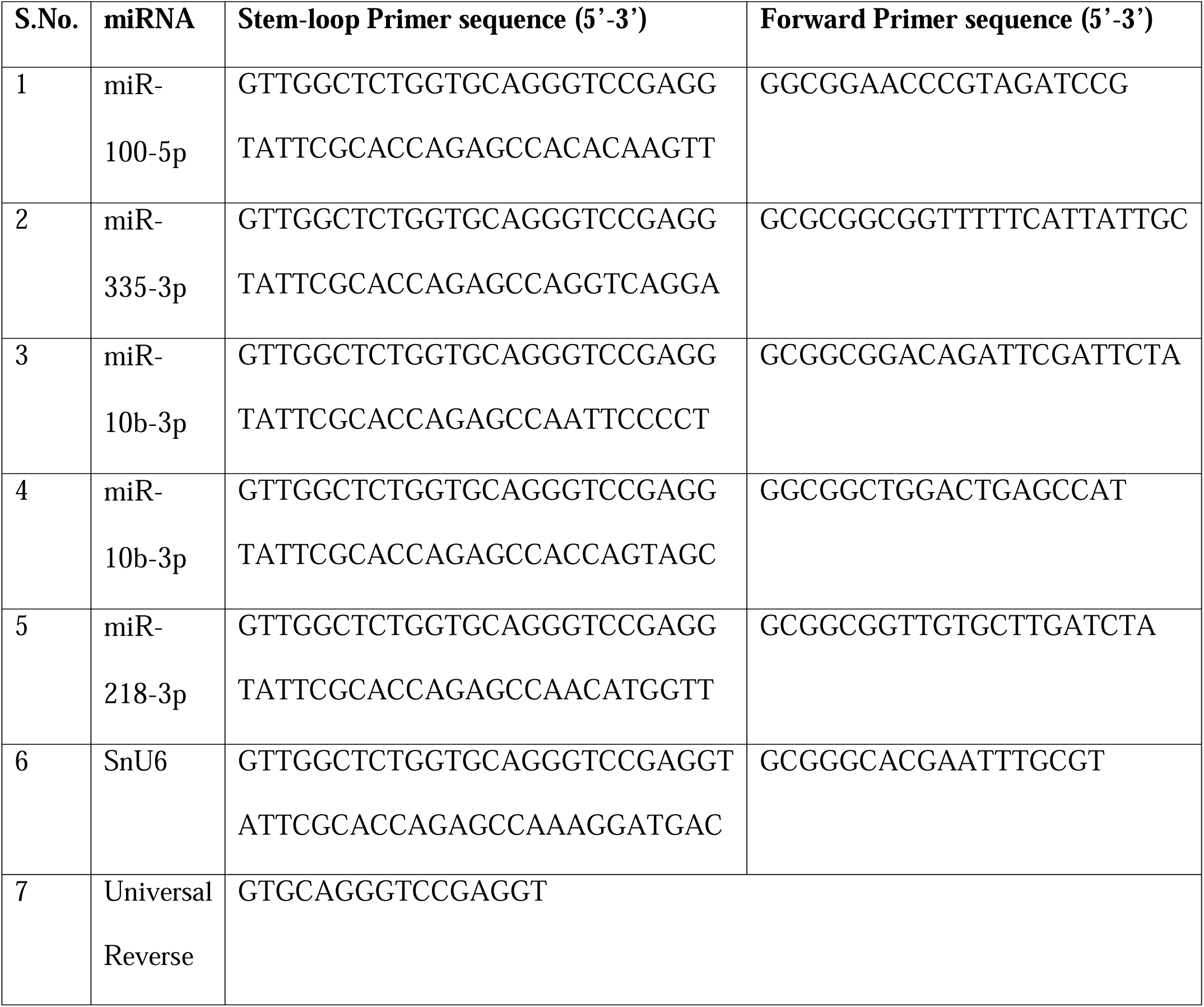
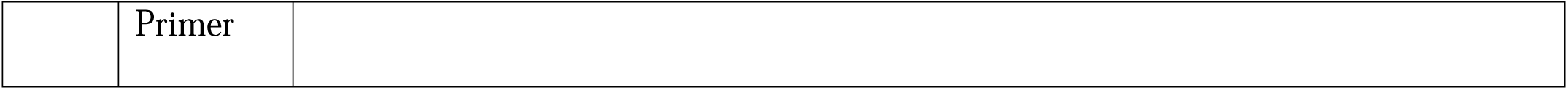
Sequences of stem-loop, forward and universal reverse primers.

### For miRNA Cluster and Targeted Gene Quantification

Transfected HEK293T and GBC cells were harvested after 48 hours of miR-17∼92 treatment. Total RNA was extracted from transfected cells and GBC tissue samples using TriZol reagent (Thermo Scientific #15596026) followed by nanodrop quantification. One step RT-qPCR was performed utilizing RNA as template using One-Step TB Green PrimeScript RT-PCR Kit II (Perfect Real Time) (Takara Bio #RR086A) according to manufacturer’s instructions using the respective primers mentioned in the Table 2.

**Table 2:**
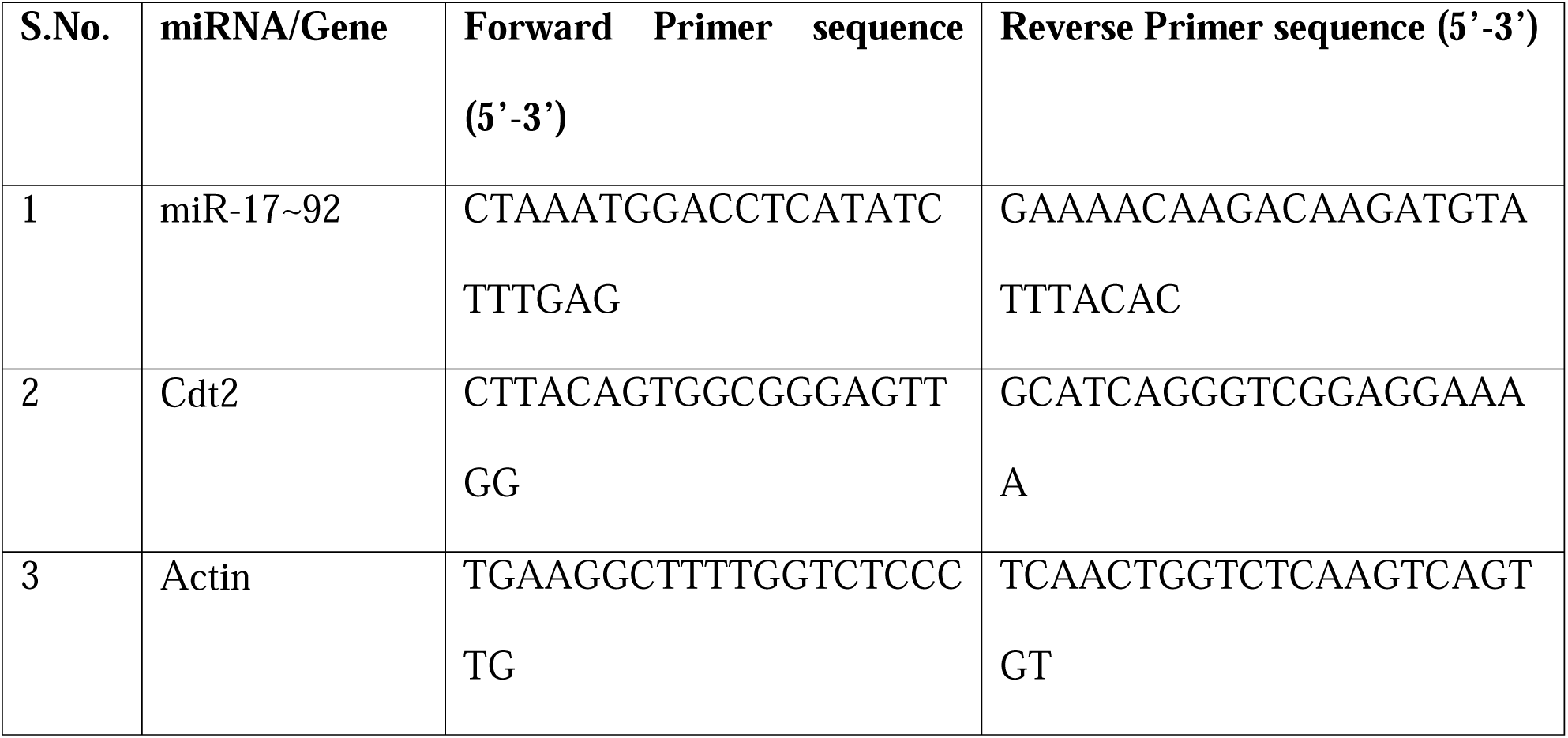
Sequences of primers.

### Western blotting

The transfected cells were harvested and total protein was extracted using radio-immunoprecipitation assay (RIPA) buffer. The extracted protein samples were resolved on SDS-PAGE (8-12%) and subjected for western blotting (12) using following antibodies: (Cdt2 (Abcam #ab72264), Actin (Invitrogen #MA5-11869), p21 (CST #2947), Set8 (CST #2996), c-Myc (CST #9402), E-cadherin (Invitrogen #701134), IgG anti-rabbit (Abcam #ab205718, IgG anti-mouse (CST #7076)). The blots were imaged using ChemiDoc system Azure Biosystems 600 (California, USA).

### Proliferation assay/growth curve

After the transfection, the cells were harvested and counted at every 24 hours of incubation (as mentioned in the result) using trypan blue dye (HiMedia #TC193-5G). The growth curve was plotted for the number of cells versus days and significance was estimated by unpaired student’s t-test.

### 3D Spheroid formation assay

Following transfection, NOZ cells were harvested and counted. A 0.7% agarose based DMEM basal layer was prepared by mixing equal volumes of 1.4% autoclaved agarose, (Genei #2600505001730) with double concentrated (2X) DMEM medium (pre-warmed at 42) and poured into 6-well culture plates. After solidification, cell suspension (10^3^ cells) was mixed with 0.7% agarose-medium at 42 (0.35% agarose-medium cell suspension). This suspension mixture was layered on the solidified agarose-medium. The plate was incubated in CO_2_ incubator overnight. Post incubation, cells were overlaid with complete DMEM medium and incubated in CO_2_ incubator (as mentioned earlier). To observe the spheroid formation, the cells were continuously monitored under phase contrast microscope for 14-16 days. At the end of the assay, the spheroids were manually counted after image acquisitions.

### Cell cycle analysis

Transfected GBC cells were harvested, pelleted down and then resuspended in 70% ethanol for overnight fixation at –20. Cells were then washed with PBS followed by resuspension of the cells in 1 ml of staining solution (100 μg/ml propidium iodide (Sigma-Aldrich #P4170-10MG), 50 mg/ml RNase-A (HiMedia #DS0003) and 0.1% Triton X-100 (SRL #30190) in PBS). The stained cells were then analysed for cell cycle progression using the CytoFLEX flow cytometer (Beckman Coulter, California, USA).

### Immunofluorescence

NOZ cells (0.8 x 10^5^) were seeded directly onto the coverslips (placed in 6 well culture plate). Following 24 hours of incubation, cells were co-transfected with miR-17∼92 or/and Flag-Cdt2 and incubated for 24 hours. Cells were then washed with PBS and fixed with 4% paraformaldehyde for 20 minutes followed by PBS wash and permeabilized with 0.1% triton X-100 (in PBS) for 20 minutes. Cells (grown on coverslip) were then blocked with 5% BSA at 4 for 1 hour, following which the cells were then incubated with primary antibody, Cdt2 (Bethyl laboratories #A300-948A) overnight in dark at 4 and then incubated with secondary antibody (Alexa flour 546 #A11010) for 1 hour in dark humid chamber (12). Lastly, the cells were incubated with DAPI 1 µg/ml (Sigma-Aldrich #D9542-1MG) for 30 minutes at 4. The excess DAPI was removed by 4-6 PBS washes. The cells were then mounted on clean slides using mounting media (Vectashield, Vector Laboratories) and observed under super resolution microscope (STED, Leica, Wetzlar, Germany).

### Annexin V apoptotic assay

Post 48 hours of transfection with miR-17∼92, NOZ cells were gently harvested and processed for annexin V apoptotic assay using “Annexin V conjugates for apoptosis detection kit” (Invitrogen, #A13199) using the manufacturer’s instructions. Briefly, the cells were washed with PBS and then resuspended in annexin binding buffer followed by incubation at RT for 10 minutes. The cells were then incubated with FITC labelled Annexin V conjugate for 15 minutes at RT. Following this, the cells were counterstained with Propidium Iodide staining solution and incubated for 5 minutes at RT in dark. The stained cells were then analysed within 30 minutes using the CytoFLEX flow cytometer (Beckman Coulter, California, USA).

### BrdU assay

miR-17∼92 transfected NOZ cells were pulsed with BrdU (30 µM) (Sigma Aldrich #B9285) for 2 hours in dark in CO_2_ incubator. The cells were then harvested and washed with PBS followed by fixation and permeabilization with 100% ethanol and triton X-100 (SRL#30190) respectively. The cells were then probed with anti-BrdU at a concentration of 0.1mg/ml (Invitrogen #B35128) for 1 hour at RT followed by incubation with secondary antibody Alexa flour 488 (Invitrogen #A-1063) for 30 minutes at RT. The cells were then washed and resuspended in propidium iodide staining solution (10 µg/ml RNase A, 100 µg/ml propidium iodide) for 30 minutes and observed using the CytoFLEX flow cytometer (Beckman Coulter, California, USA).

### Wound healing assay

Equal numbers of miR-17∼92 treated and control GBC cells were seeded into 24 well plate and incubated for 24 hours in the CO_2_ incubator. After attaining a confluency of 85-90%, a wound was created with the help of 200 µl micropipette tip. The cells were observed and assessed for the wound healing or migration under phase contrast microscope till the wound closure.

### Invasion assay

miR-17∼92 transfected GBC cells (0.8×10^5^) were resuspended in a serum-free DMEM. Further, 2 mg/ml ECM gel (Sigma-Aldrich #e1270) was coated on the upper chamber (insert) of transwell plate (Corning, New York, USA) and allowed to solidify at 37 for 2 hours. The cells suspensions were seeded on ECM layer in insert (upper chamber). In the lower chamber, DMEM supplemented with 10% FBS was added. The cells were grown for 48 hours as mentioned earlier. After incubation, invaded cells (on the outer surface of insert) were fixed with 5% glutaraldehyde (Sigma-Aldrich #G6257). The cells were then stained with 2% crystal violet solution (SRL #28376; in 2% v/v ethanol), observed and counted under inverted microscope.

### Statistical Analysis

All the western experiments and RT-qPCR were performed in biological triplicates and duplicates respectively. The growth curve, wound healing and invasion assays were performed in biological/experimental triplicates as well. The data are presented as the mean ± S.E. and unpaired student’s t-test was performed to calculate the significant p-value. ImageJ software was used to quantify the intensity of protein bands, fluorescence intensity and wound width of cells wherever applicable.

## Results

### miRNA sequencing showed significant differential expression of 388 miRNAs in GBC cells

Quality assured RNA from NOZ and HEK293T cells (RIN score of 9.3) were used for small RNA extraction followed by next-generation sequencing to prepare the miRNA landscape in GBC. A total of 14,265,080 and 13,629,498 trimmed reads were obtained for HEK293T and NOZ respectively, which were mapped back to the reference genome. The mapped reads of HEK293T and NOZ were then graphically arranged to demonstrate the reads on each chromosome (Figure 1A). Some of the mapped reads were found overlapped with multiple regions, thus each sRNA read was annotated to unique annotation preferentially in the order of known miRNA > rRNA > tRNA > snRNA > snoRNA > repeat > gene > novel miRNA (Figure S1). Mapped and annotated reads were then analyzed with specific sequences in the miRBase to procure detailed information of the miRNAs. The analysis resulted into the identification of 1220 already annotated miRNAs and discovery of 37 novel miRNAs. The expressions of these 1257 miRNAs in each sample were statistically analyzed and normalized to build a TPM density distribution graph which represents the miRNA expression variation between the two samples (Figure 1B). The Pearson correlation coefficient (R^2^ = 0.657) between the two cell types shows a moderate yet significant positive correlation of miRNA expression (Figure 1C; Figure S2). Among these miRNAs, 259 were primarily present in HEK293T cells while 281 were exclusively expressed in NOZ cells and 717 miRNAs were differentially expressed in both HEK293T and NOZ cells. Out of 717 DEmiRNAs, 388 shows significant differential expression profile between HEK293T and NOZ cells (Figure 1D; Supplementary file 1). Specifically, 191 miRNAs were upregulated and 197 miRNAs were downregulated in NOZ cells relative to HEK293T cells (Figure 1E, Supplementary file 1), as visualized in the hierarchical clustering heat map (Figure 1F, Supplementary file 2).

**Figure 1:**
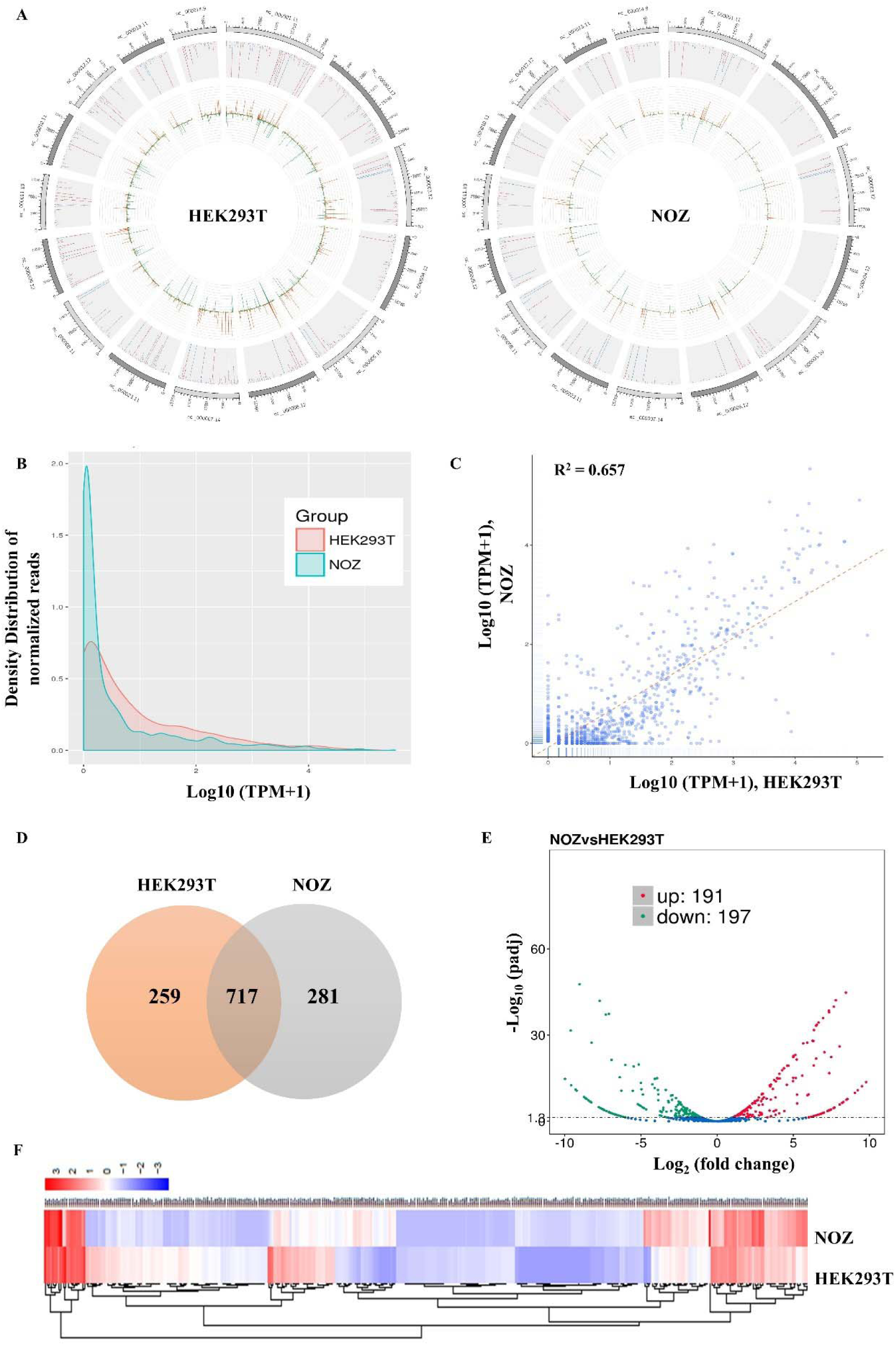
Analysis of differentially expressed miRNAs. **A.** Reads distribution per chromosome in HEK293T and NOZ cells. 10 longest contigs were plotted**. B.** TPM density distribution graph depicting the expression of normalized reads for miRNAs in each sample. x-axis represents the log_10_ value for normalized reads of miRNAs; y-axis represents density distribution of the normalized reads. **C.** Pearson correlation between HEK293T and NOZ cells revealing that the two samples are positively correlated to each other with Pearson correlation coefficient (R^2^). x-axis represents log_10_ value for normalized reads of HEK293T miRNAs; y-axis represents log_10_ value for normalized reads of NOZ miRNAs; Each data point represents the expression of a single miRNA in each cell type. **D.** Venn diagram of differentially expressed miRNAs in HEK293T and NOZ cells showing 259 miRNAs expressed exclusively in HEK293T, 281 miRNAs in NOZ and 717 miRNAs differentially expressed in both the cells. **E.** Volcano plot showing 191 upregulated and 197 downregulated miRNAs in NOZ cells. x-axis represents log_2_ fold change, y-axis represents statistically significant difference in expression where padj is adjusted p-value or q-value (threshold<0.05); Dots represent each miRNA, blue dots indicate no significant difference, red dots indicate significant upregulation, green dots indicate significant downregulation. **F.** Heat map showing varying expression level of each miRNA in both the cell lines where red indicates the highest and blue indicates the lowest expressed miRNAs.

### miRNA landscape discovered 37 novel miRNA signatures

From the sequencing data, novel miRNAs were discovered by analyzing the characteristic hairpin structures of miRNA precursors using miREvo tool and miRDeep2 software. The analysis led to the discovery of 37 novel miRNAs (which were not annotated previously). Among these, 19 novel miRNAs were differentially expressed in both the cell types (HEK293T and NOZ), while 10 novel miRNAs were only present in non-cancerous cells (HEK293T) and 8 were exclusive to GBC cells (NOZ) (Supplementary Table 2; Supplementary file 3). This novel miRNA expression profile uncovers 27 novel miRNAs (including both 19 DE and 8 exclusives to GBC) expressed in GBC cells strengthening them as signature novel miRNAs profile for GBC. Upon further analysis, we found that miRNAs: novel_107, novel_154, novel_161, novel_218, novel_222, novel_289, novel_309 and novel_321 shows significant differential expression. miRNAs novel_107, novel_161, novel_218, novel_222, novel_289, novel_309 and novel_321 being downregulated (q-value ≤ 0.05 and Log_2_ FC ≤ –1) and miRNA novel_154 being upregulated (q-value ≤ 0.05 and Log_2_ FC ≥ 1) in NOZ cells. To elucidate the potential biological functions of eight GBC exclusive novel miRNAs, we performed target prediction using established miRNA target prediction algorithms. Notably, the predicted targets included genes implicated in cell cycle regulation (e.g., CDK4, CCNE1, E2F1), apoptosis (BAX, BCL2L1), and key signaling pathways (AKT1, MAPKAPK3, STAT6), as well as genes involved in immune regulation (PDCD1, CIITA, NFKBIB, CXCL5) (Figure 2A, Supplementary file 6). Our lab is engaged methodically in the nomenclature and validation of these miRNAs to facilitate their functional and mechanistic characterization.

**Figure 2:**
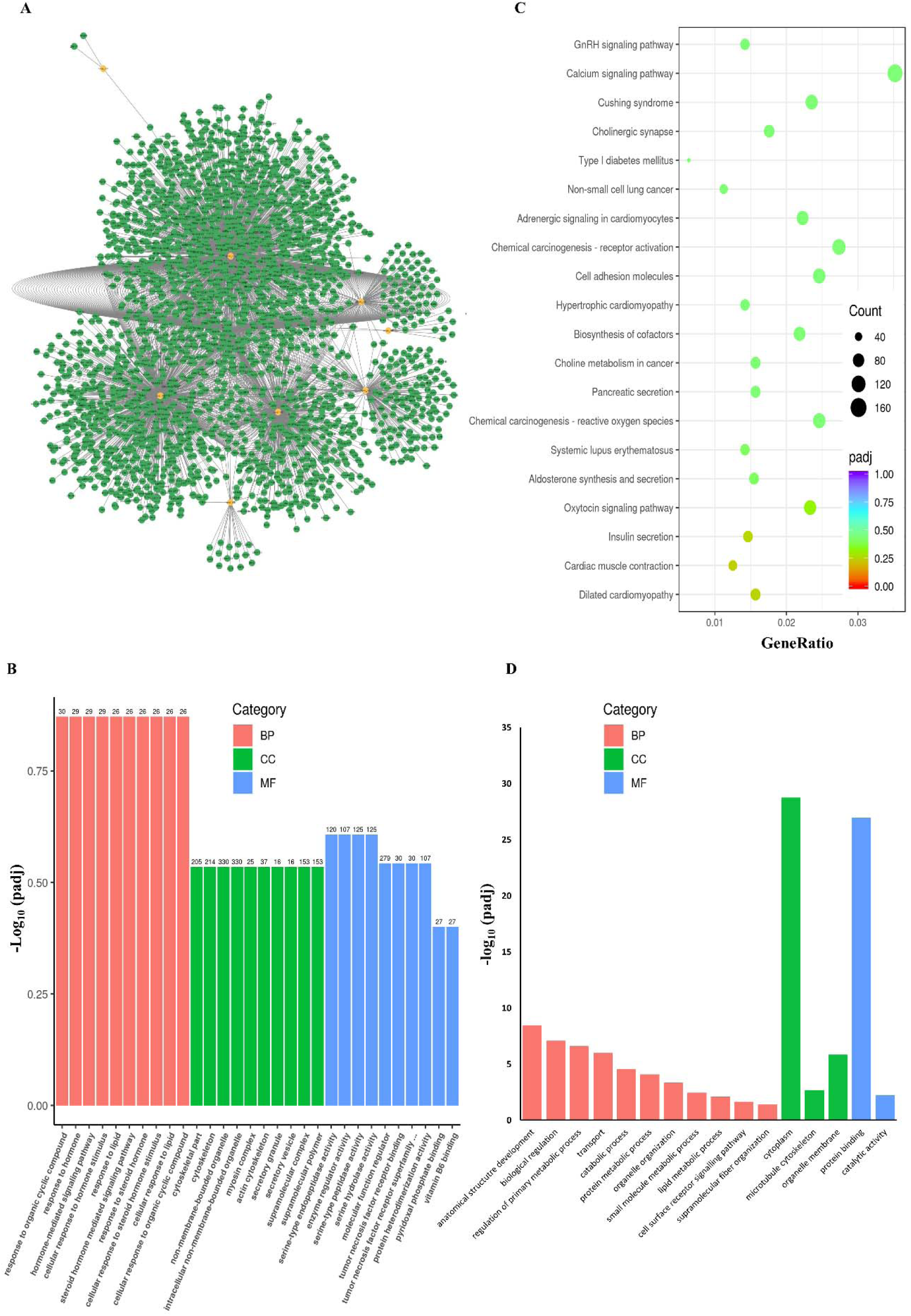
Network and Pathway enrichment analysis. **A.** miRNA-mRNA network interactome between GBC exclusive novel miRNAs and their predicted mRNA targets. The source nodes denoted with orange colour represent the novel miRNA while the green nodes represent the predicted mRNA targets. **B.** GO analysis of DE miRNAs in NOZ cells showing top 10 Biological Processes (BP), 10 Cellular Components (CC) and 10 Molecular Function (MF). On the y-axis, –log_10_ (padj) represents –log_10_ of corrected/adjusted p-value in hypergenometric test; Numbers on the bar represent the count of target genes in the related process. **C.** KEGG enrichment analysis of predicted targets of DE miRNAs in NOZ cells enlisting top 20 pathways. On the x-axis, GeneRatio represents the ratio of target genes to total genes in KEGG enrichment; Count represents number of target genes related to the particular pathway; padj represents the corrected/adjusted p-value in hypergenometric test. **D.** GO analysis of 8 exclusively expressed novel miRNAs in GBC cells most enriched Biological Processes (BP), Cellular Components (CC) and Molecular Function (MF).

### Target prediction and pathway enrichment of differentially expressed miRNAs

To have a closer look on the targets and pathways regulated by these differentially expressed (DE) miRNAs, we performed the miRNA-target prediction hunt on RNAhybrid and miRanda. We then performed the enrichment analysis on the predicted targets of DE miRNAs for Gene Ontology (GO) function and Kyoto Encyclopedia of Genes and Genomes (KEGG) pathway. Functional annotation of DE miRNAs by GO analysis showed enrichment of potential target genes in cellular processes like tumor necrosis, hormone mediated signaling pathways, cellular responses to lipids and steroids etc. (Figure 2B; Supplementary file 4). The KEGG analysis of these differentially expressed (DE) miRNAs revealed significant enrichment in pathways such as cell adhesion molecules, apoptosis, ferroptosis, pathways in cancer, gastric cancer, microRNAs in cancer, the p53 signaling pathway, and the calcium signaling pathway, among others. Additionally, it was observed that these DE miRNAs are implicated in pathways that regulate oncogenesis, chemical carcinogenesis, cell adhesion, cell cycle, and apoptosis in gallbladder cancer cells (Figure 2C; Supplementary file 5).

Furthermore, the targets of eight GBC exclusive novel miRNAs are significantly enriched in molecular functions like protein binding (padj < 10^-25^) and catalytic activity suggesting their involvement in protein-protein interactions and enzymatic processes in GBC. Enriched biological processes included anatomical structure development, biological regulation, regulation of primary metabolic process, transport, and various metabolic and catabolic processes.

Notably, cell surface receptor signaling pathway and supramolecular fiber organization were also represented, indicating roles in cell signaling and cytoskeletal dynamics. The most enriched cellular component was the cytoplasm (padj <10^-27^), followed by organelle membrane and microtubule cytoskeleton, highlighting the potential involvement of these miRNAs in regulating cytoplasmic processes, membrane dynamics, and cytoskeletal organization (Figure 2D).

### Validation of differentially expressed miRNAs in GBC tissue and cell lines

After profiling the miRNA expression landscape in GBC cells, we proceeded to evaluate the reliability of our findings by validating the expression levels of the four most significantly differentially expressed (both upregulated and downregulated) miRNAs in GBC cell lines. Among the selected candidates, the relative expression of miR-100-5p and miR-335-3p was higher in GBC cell lines, NOZ and G415 in comparison to HEK293T cells (Figures 3A-B). Whereas, the levels of miR-10b-3p and miR-218-3p were downregulated in GBC cells (Figures 3C-D). The forementioned miRNA levels were also validated in clinical tissue specimens obtained from GBC patients to confirm and assure the biological relevance of the identified miRNA signatures. GBC tissues exhibited significantly higher levels of miR-100-5p and miR-335-3p than non-cancerous gallbladder tissues (gallstone samples; Figures 3A-B). On the other hand, GBC tissues showed a considerable downregulation of miR-10b-3p and miR-218-3p (Figures 3C-D), corroborating the findings in the cell line models.

**Figure 3:**
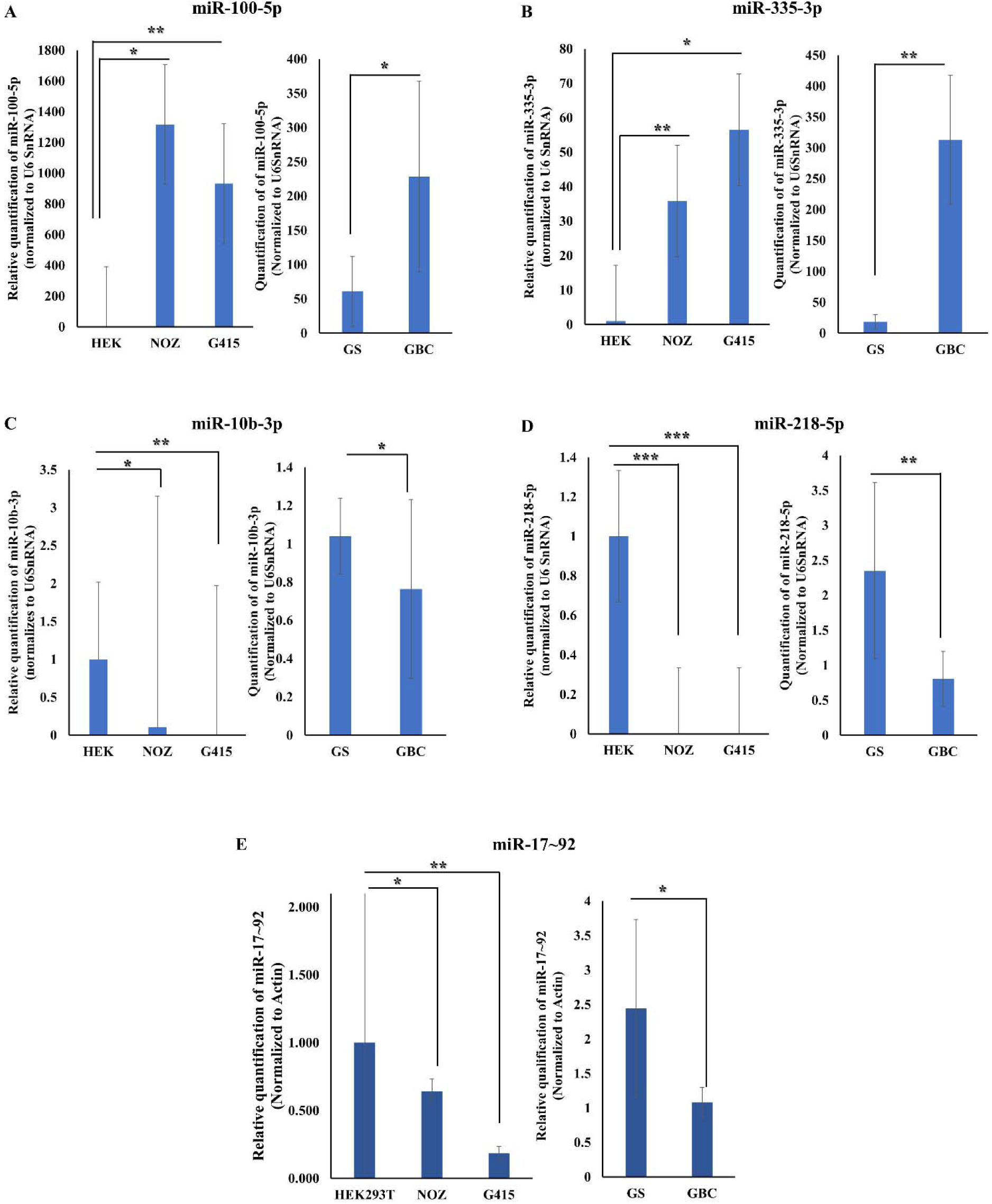
Validation of DE miRNAs. **A.** qRT-PCR analysis of up-regulated miRNA miR-100-5p levels in GBC cells and GBC patient tissue (n=5) vs GS (n= 5). **B**. qRT-PCR analysis of up-regulated miRNA miR-335-3p levels in GBC cells and GBC patient tissue (n=5) vs GS (n= 5). **C**. qRT-PCR analysis of down-regulated miRNA miR-10b-3p levels in GBC cells and GBC patient tissue (n=3) vs GS (n= 3). **D**. qRT-PCR analysis of down-regulated miRNA miR-218-5p levels in GBC cells and GBC patient tissue (n=5) vs GS (n= 5). **E.** qRT-PCR analysis of miR-17∼92 levels in GBC cells and GBC patient tissue (n=3) vs GS (n= 3). Error bars represent standard error (SE). * Represents p value < 0.05, ** represents p value < 0.01 & *** represents p value < 0.005.

### miRNA sequencing revealed miR-17∼92 to be downregulated in GBC cells

From the sequencing data analysis, we found that among DE miRNAs, miR-17-5p, miR-18a, miR-19a-3p, miR-19b-1-5p, miR-20a and miR-92a-1-5p were downregulated in gallbladder cancer cell line, NOZ (Supplementary file 1). These miRNAs together form a miRNA cluster known as miR-17∼92 (18). To validate the above result, we checked the expression of miR-17∼92 cluster in GBC cells as well as in tissue samples from GBC patients, confirming that its expression is significantly suppressed in GBC cells and patient tissue samples (Figure 3E).

### Ectopic expression of miR-17∼92 suppresses Cdt2 expression in GBC cells

Previous literatures have shown that Cdt2/DTL is highly amplified in many cancers, including gallbladder cancer (GBC cell line SNU308, The Human Protein Atlas (TCGA) database). We also evaluated the relative expression of Cdt2 transcript in gallbladder cancer patient tissues and found it to be significantly upregulated (Figure 4A). Next, we performed the miRNA database hunt to check the predicted targets of miR-17∼92 and we found that various miRNAs of the cluster directly target the 3’UTR of Cdt2 transcript. Therefore, we aimed to investigate whether miR-17∼92 targets Cdt2 and also to determine the potential implications of this interaction. To achieve the same, we transfected HEK293T and GBC cells with miR-17∼92 and found that Cdt2 protein is significantly suppressed in GBC cells upon ectopic expression of miR-17∼92 both at transcript as well as at protein level, while there is no significant effect of miR-17∼92 on Cdt2 expression in HEK293T cells (Figures 4B, 4C and 4D). The above finding confirmed that miR-17∼92 directly suppressed Cdt2 in GBC cells (Figures 4B-D). After establishing the relation between miR-17∼92 and Cdt2, we wanted to check how miR-17∼92 regulate the expression of the downstream targets of Cdt2 (especially the cell cycle regulators p21 and Set8, which are downstream targets of CRL4^Cdt2^ complex) and found that suppression of Cdt2 by miR-17∼92 leads to stabilization of S-phase licensing factor Set8 in GBC cells (Figures 4E and 4F; Figure S3). But surprisingly, we noticed that the level of p21 protein was also suppressed (Figures 4E and 4F). Further, we checked the status of c-Myc (oncoprotein) in GBC cells as c-Myc and miR-17∼92 work in a negative feedback loop mechanism. We confirmed that c-Myc is significantly reduced in GBC cells upon miR-17∼92 transfection (Figures 4E and 4F; Figure S3). Since, Cdt2 is a S-phase regulatory protein and is remain localized to the nucleus during S-phase of cell cycle. Therefore, we want to check whether the suppression of Cdt2 by miR-17∼92 has any effect on the localization of the Cdt2 protein. We performed super resolution microscopy and found that the Cdt2 is highly enriched in the nucleus of NOZ cells (control) (Figure 5A; upper panel; Figure 5B) but ectopic expression of miR-17∼92 resulted in the localization of Cdt2 in the cytoplasm with an overall reduced intensity of Cdt2 (Figure 5A; middle panel; Figure 5B). The overexpression of miR-17∼92 has also resulted into distortion of the cellular morphology of NOZ cells and expulsion of some of the nuclear content (Figure 5A; middle panel). While the co-transfection of miR-17∼92 along with Flag-Cdt2 has rescued the expression of Cdt2 which is more prominent in the nucleus (Figure 5A; lower panel; Figure 5B). The co-transfection of Flag-Cdt2 with miR-17∼92 has also restored the cellular morphology of the cells (Figure 5A; lower panel).

**Figure 4:**
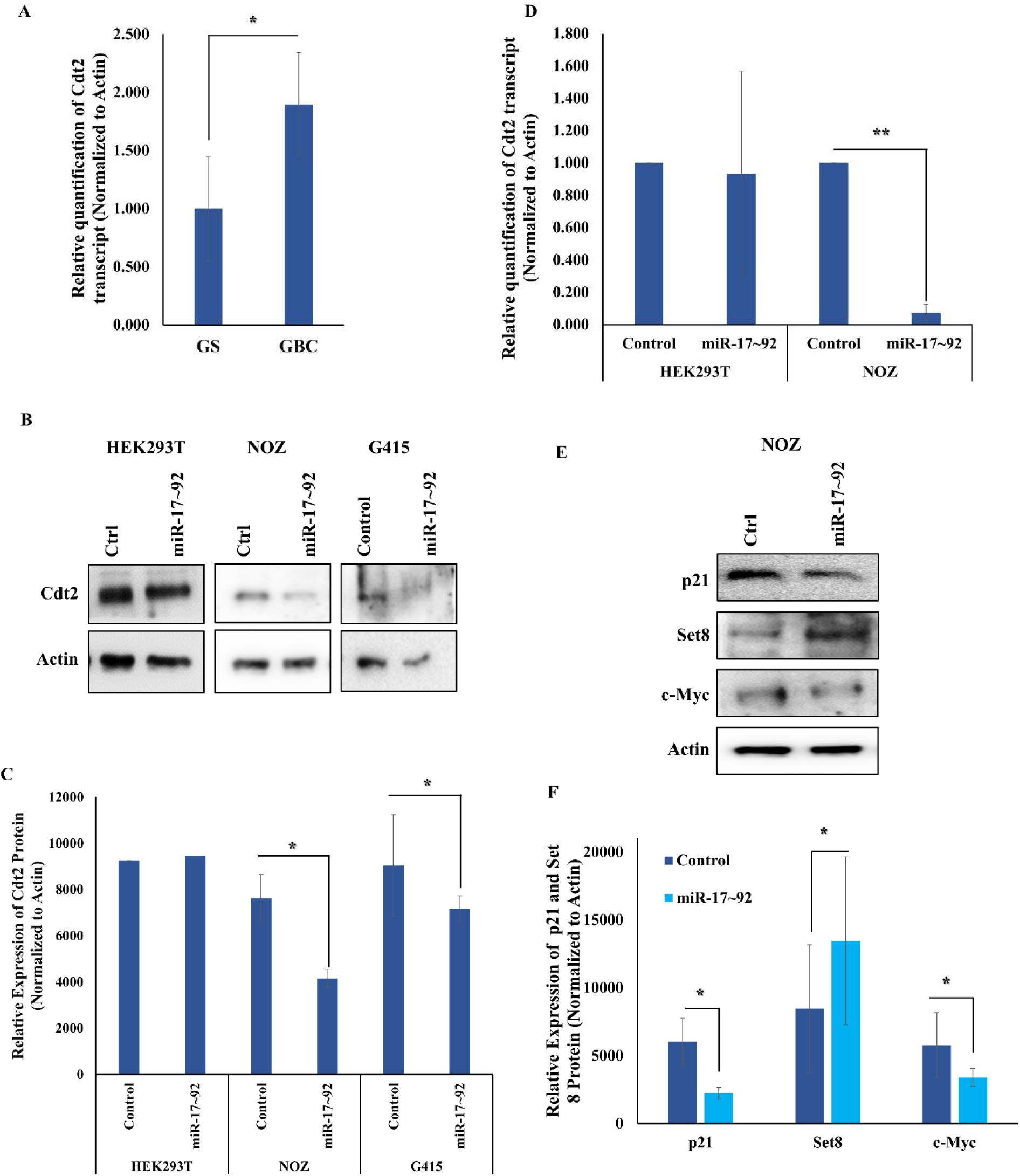
Ectopic expression of miR-17∼92 destabilizes Cdt2. **A.** qRT-PCR analysis of Cdt2 transcript level in GS (n= 4) and GBC (n= 4) tissue samples. **B.** Western blot analysis for Cdt2 protein shows suppression in Cdt level in miR-17∼92 treated GBC cells while no effect in HEK293T cells. **C.** Quantification of Cdt2 protein level analysis in HEK293T and GBC cells after miR-17∼92 transfection. **D.** qRT-PCR analysis showing miR-17∼92 suppresses the Cdt2 transcript levels in NOZ cells only. **E.** Western blot analysis for p21, Set8 and c-Myc proteins results in increased level of Set8 and suppressed levels of p21and c-Myc in miR-17∼92 treated NOZ cells. **F.** Quantification of p21, Set8 and c-Myc levels analysis in NOZ cells after miR-17∼92 transfection. Error bars depict S.E. * represents p-value<0.05. *** represent p-value<0.001.

**Figure 5:**
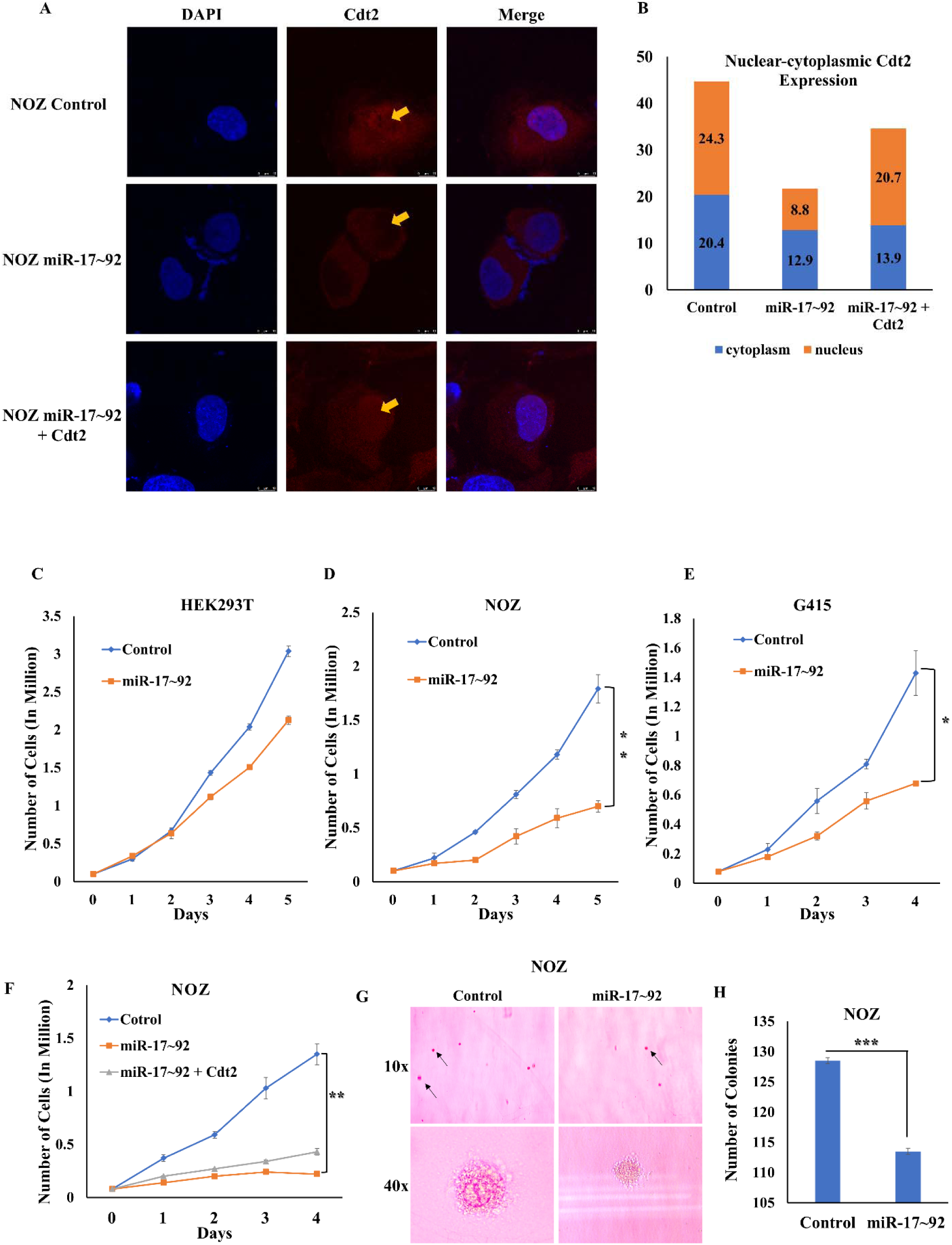
Ectopic expression of miR-17∼92 suppresses proliferation of GBC cells. **A.** NOZ cells were transfected with miR-17∼92 and miR-17∼92 + Cdt2 cocktail for 48 hours, incubated with anti-Cdt2 and analyzed under confocal microscopy (yellow arrow represents nuclear localization of Cdt2 protein). Scale bar-10 μM. **B.** Stacked column graphical representation of nuclear-cytoplasmic content of Cdt2 protein. Results are shown as mean of 5 different randomly selected cytoplasmic and nuclear regions (same area) and Cdt2 intensity levels were quantified. **C.** Growth curve of HEK293T cells after ectopic expression of miR-17∼92 showing no significant change. **D.** Growth curve of NOZ cells shows significant reduction in proliferation after ectopic expression of miR-17∼92. **E.** Growth curve of G415 cells shows significant reduction in proliferation after ectopic expression of miR-17∼92. **F.** Rescue-growth curve of NOZ cells co-transfected with miR-17∼92 and Flag-Cdt2 along with respective controls. **G.** 3D spheroid formation assay showing the effect of miR-17∼92 on the spheroid formation ability of NOZ cells at 10x and 40x (black arrow represents spheroids). **H.** Graphical representation of miR-17∼92 suppresses the colony formation ability of NOZ cells. Error bars depict S.E. ** represents p-value<0.01; *** represents p-value<0.001.

### miR-17∼92 Inhibits the proliferation of GBC cells

After establishing that miR-17∼92 destabilizes Cdt2 level in GBC cells, next, we wanted to check the effect of miR-17∼92 on the growth and proliferation of the GBC cells. We observed that the ectopic expression of miR-17∼92 suppresses the growth and proliferation of both NOZ and G415 cells (Figures 5D and 5E) while no significant effect was observed on HEK293T cells (Figure 5C). Also, miR-17∼92 expression caused morphological changes including distorted cellular structure in GBC cells while no such change was observed in the HEK293T cells (Figure S4). Further, we checked if the suppressed proliferation rate of the GBC cells, due to miR-17∼92 expression, could be rescued by ectopic expression of Cdt2. To confirm this, we transfected NOZ cells with either miR-17∼92 alone or co-transfected with miR-17∼92 and Flag-Cdt2. We found that co-transfection of miR-17∼92 with Cdt2 could rescue the proliferation of NOZ cells only to some extent (Figure 5F). We also performed spheroid formation assay to check the effect of miR-17∼92 on ability of GBC cells to form 3D spheroids and observed that ectopic expression of miR-17∼92 in NOZ cells significantly reduced both the number/colonies as well as size of spheroids (Figures 5G and 5H).

### miR-17∼92 expression arrests GBC cells in the S-phase of the cell cycle and induces apoptosis in GBC cells

Since we observed that miR-17∼92 suppresses the expression of major cell cycle regulator Cdt2 which in turn checks proliferation of these cells, we wanted to see the effect of miR-17∼92 on different phases of the cell cycle. For the same, we analyzed the cell cycle and found that the ectopic expression of miR-17∼92 arrested the GBC cells in S-phase of the cell cycle (Figures 6A and 6B, Figure S5; control vs miR-17∼92 treated). We have also found significant increase in the sub-G1 population representing dead cells with fragmented DNA which could either be apoptotic or necrotic population (NOZ cells treated with miR-17∼92, Figure 6A vs 6B). To validate this observation, we performed annexin V apoptotic assay in the NOZ cells. The results revealed a notable increase in the both early and late apoptotic population from 8.11% and 2.04% to 13.57% and 13.87% respectively with an overall increase of 17.29% in apoptotic rate upon miR-17∼92 transfection in NOZ cells (Figure 6C and D). The ectopic expression of miR-17∼92 has also induced a small amount of necrosis from 6.71% to 12.43% (Figure 6C and 6D).

**Figure 6:**
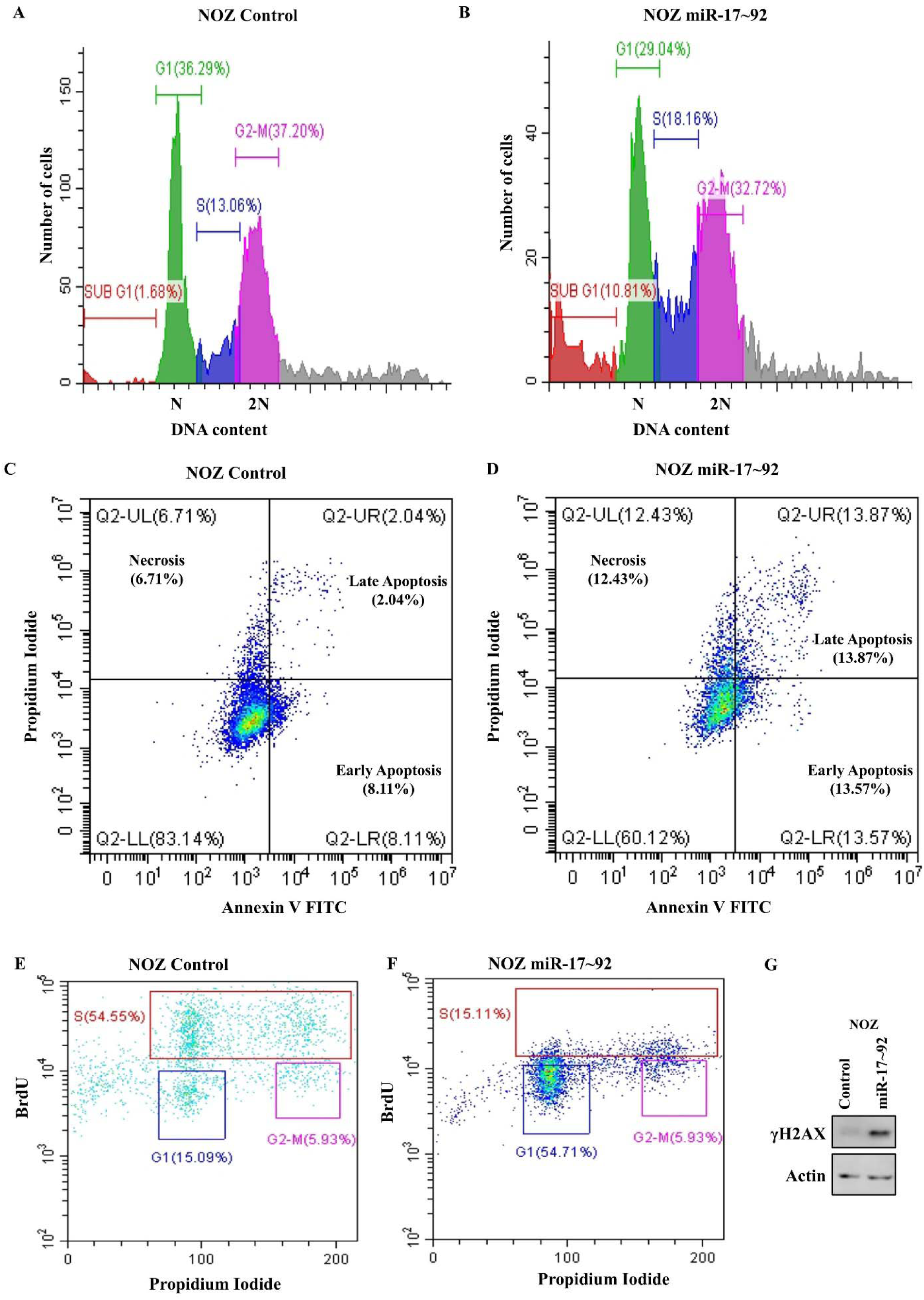
Ectopic expression of miR-17∼92 induces S-phase cell cycle arrest and apoptosis in GBC cells. **A.** Flow cytometric analysis of Cell cycle in NOZ control cells. **B.** Flow cytometric analysis of Cell cycle representing miR-17∼92 induces S-phase arrest and increased sub G1 population in miR-17∼92 treated NOZ cells indicating fragmented DNA resulted from apoptosis/necrosis. **C.** Annexin V detection with flow cytometric analysis in NOZ control cells **D.** Annexin V detection with flow cytometric analysis demonstrating miR-17∼92 induces apoptosis as well as necrosis in NOZ cells. **E.** BrdU incorporation and flow cytometric analysis in NOZ control cells **F.** BrdU incorporation and flow cytometric analysis showing decreased BrdU incorporation in Cdt2 deprived NOZ cells. G. Western blot analysis for γH2AX proteins results in increased level of γH2AX in miR-17∼92 treated NOZ cells.

Since Cdt2 is a replication licensing factor and earlier we observed that suppression of Cdt2 by miR-17∼92 resulted in S-phase arrest, therefore we further investigated the status of DNA synthesis upon miR-17∼92 in GBC cells. To understand the same we performed BrdU incorporation assay. The results revealed significantly reduced incorporation of BrdU from 54.55% to 15.11% upon miR-17∼92 treatment (Figure 6E and 6F) in NOZ cells, which confirms decrease replication rate of the treated cells. To further explore the replication/DNA damage status, we checked the phosphorylation status of H2AX and found an increase in H2AX phosphorylation upon miR-17∼92 expression indicating increase in replication and DNA damage stress (Figure 6G).

### miR-17∼92 also inhibits the metastatic capabilities of GBC cells

Since miR-17∼92 suppresses the growth and proliferation of GBC cells, we next wanted to see whether it can affect the metastatic abilities of these cells. Upon performing wound healing and invasion assays, we observed a significant suppression in the wound healing or the migratory ability of GBC cells (Figures 7A and 7B) after miR17-92 expression. We also found that miR-17∼92 expression significantly reduced the invasive ability of NOZ cells (by ∼28%) and G415 cells (by ∼31%, Figures 7C and 7D). Since epithelial to mesenchymal transition (EMT) is a major phenomenon involved in the metastatic process of cancerous cells, therefore, we checked the level of EMT biomarker, E-cadherin (epithelial cell marker) and found that miR-17∼92 expression significantly increased the level of E-cadherin in NOZ cells, suppressing its metastatic behavior (Figures 7E and 7F).

**Figure 7:**
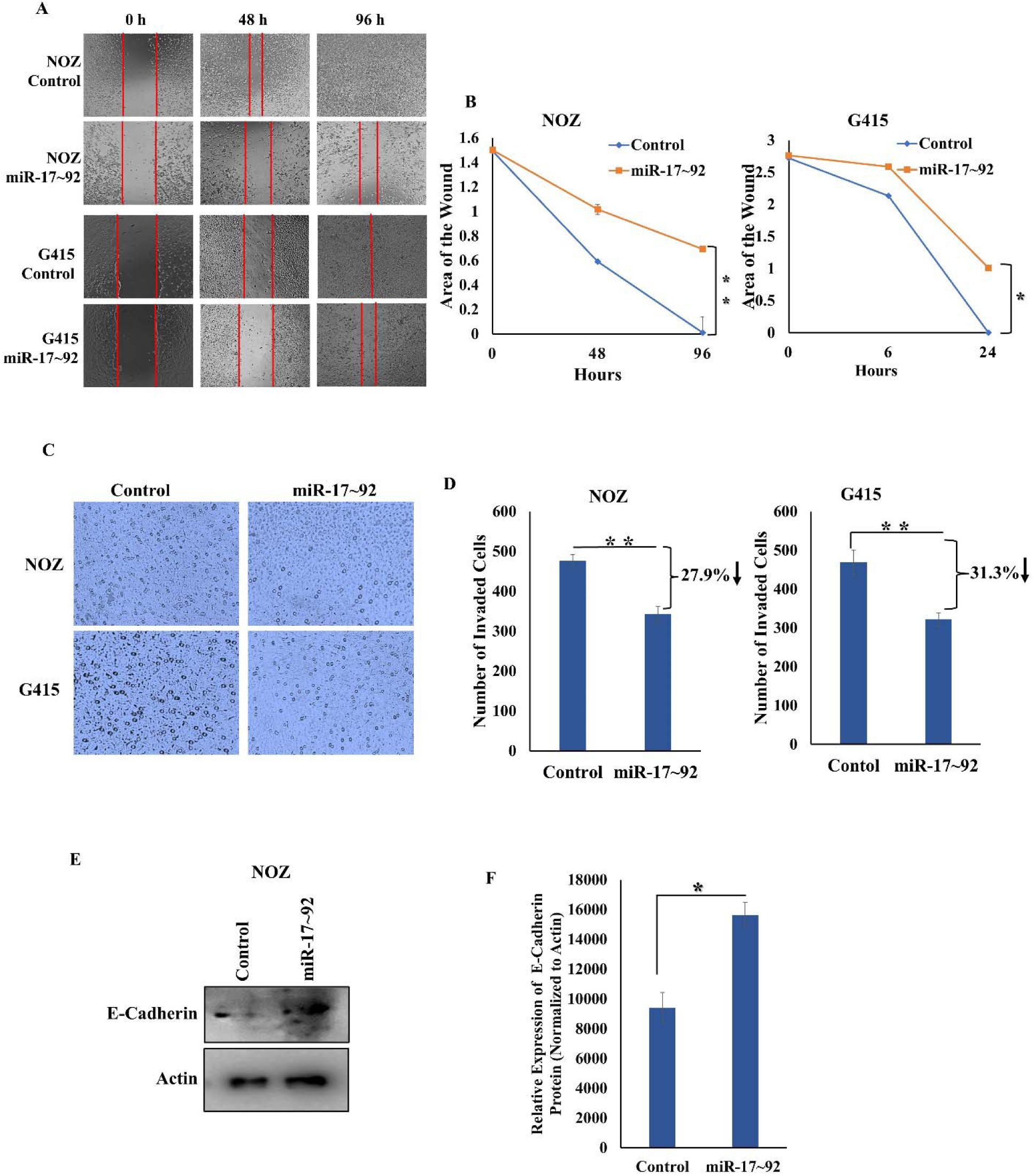
Ectopic expression of miR-17∼92 suppresses the metastatic abilities of GBC cells. **A.** Phase contrast microscopic images of control and miR-17∼92 treated NOZ and G415 cells at different time interval representing reduction in the migratory ability of treated cells. **B.** Graphical representation of wound healing assay of NOZ and G415 cells with respective control. **C.** Phase contrast microscopic images of control and miR-17∼92 treated NOZ and G415 cells representing invasive ability of treated cells. **D.** Graphical representation of invasion assay of GBC cells with respective control, showing miR-17∼92 suppresses the invasiveness of NOZ cells by 28% and G415 cells by ∼31%. **E.** Western blot analysis showing miR-17∼92 increases the E-cadherin level in NOZ cells. **F.** Quantification of E-Cadherin level in NOZ cells after miR-17∼92 transfection results in the increased expression of E-cadherin level. 8 different areas were selected for the transwell assays and number of cells were counted. Error bars depict S.E. * represents p-value<0.05; ** represents p-value<0.01.

## Discussion

MicroRNAs play very important regulatory role in various molecular and cellular processes. The dysregulation of which is associated with the onset and progression of various pathological conditions including cancers (15). Despite being a key regulator of gene expression, the interplay between the miRNAs and GBC progression is minimally explored. To understand the role of miRNAs in GBC oncogenesis, the study establishes a foundational atlas of miRNA expression in GBC and emphasizes the importance of integrative validation strategies.

By integrating miRNA sequencing with functional validation, this study delineates the miRNA landscape in GBC, identifying 1,220 known miRNAs and discovering 37 novel miRNAs (Pearson correlation coefficient >0.5), of which 388 miRNAs exhibited significant differential expression (191 upregulated, 197 downregulated) between NOZ and HEK293T cells (Figure 1C, E). This extensive profiling underscores the intricate regulatory networks orchestrated by miRNAs in GBC, consistent with the emerging view that miRNAs are central to cancer pathophysiology. These miRNAs accumulate to concentrations critical for target regulation, with select species (e.g., miR-155-5p, miR-100-5p) exceeding 9,000 reads per million (RPM) in GBC cells-a threshold indicative of biological relevance in oncogenic pathways. Notably, miR-155-5p has previously been shown to enhance the proliferation and invasion of GBC cells, reinforcing its tumorigenic potential (19,20). Conversely, miR-218-5p exhibited the most pronounced downregulation (log2 fold change = –11.2, *q* < 0.001; Supplementary file 1, Figure 3D), which is sponged by a long non-coding RNA, CCAT1 to upregulate the oncogenic protein Bmi-1, has been implicated in promoting GBC progression (20). Importantly, miR-218-5p has also been shown to sensitize GBC cells to gemcitabine, suggesting a role in enhancing chemotherapy response (21). This positions miR-218-5p as a promising candidate for targeted therapy. Apart from miR-218-5p our validation in patient-derived tissues, shows miR-100-5p and miR-335-3p were upregulated, and miR-10b-3p was downregulated, thus reinforcing the biological relevance of these miRNAs as potential diagnostic or prognostic biomarkers.

Recent advances in liquid biopsy have highlighted the diagnostic utility of circulating miRNAs. In line with this, our findings of elevated miR-552-3p and miR-372-3p in GBC cells mirror those reported by Yang et al. (2022), who demonstrated that these miRNAs are secreted into the circulation and may facilitate tumor microenvironment remodeling and metastasis (22,23). The convergence of tissue and plasma data underscores the robustness of these miRNAs as diagnostic biomarkers, particularly when combined with established markers such as CEA and CA19-9.

Additionally, the novel miRNAs discovered in GBC appear to regulate a wide spectrum of genes involved in protein interactions, metabolism, signaling, cell cycle, apoptosis, immune response, and cytoskeletal organization, highlighting the multifaceted roles these novel miRNAs may play in gallbladder cancer biology (Figure 2A and 2D). The findings underscore the complex regulatory networks orchestrated by these novel miRNAs in GBC and warrant further investigation into their specific roles serve as potential biomarkers or therapeutic targets in GBC.

Here, we also reported dysregulation of various miRNA clusters such as miR-23a/27a/24-2, miR-106a∼363, miR-99b/let-7e/miR-125a, miR-199a/214, etc. were upregulated while miR-99a/let-7c, miR-143/145, miR-194/215, etc. were downregulated. Interestingly, among all the clusters, miR-17∼92 cluster was significantly downregulated in GBC cells as well as in GBC patients’ tissues (Figure 3E). The cluster has been in discussion for its contradictory differential role across different sets of cancer (24). We also show that in GBC, miR-17∼92 exhibit a tumor suppressive role by targeting an essential yet oncogenic cell cycle factor Cdt2 at its transcript level (Figure 4D), leading to its reduced protein (Figures 4B and 4C). Suppression of Cdt2 leads to stabilization of Set8 protein (a licensing factor; Figures 4E and 4F). The suppression of CRL4^Cdt2^ was also expected to stabilize the level of p21 protein but, p21 is one of the direct targets of miR-17∼92, thus, it destabilizes p21 protein (Figures 4E and 4F). This observation establishes the proof of concept (25). Aligning with the above findings, our confocal microscopy analysis demonstrated a significant reduction in both nuclear as well as cytoplasmic level of Cdt2 following miR-17∼92 treatment. This decrease in Cdt2 signal intensity infers that miR-17∼92 likely regulate the Cdt2 at post transcriptional level and newly synthesized Cdt2 is insufficient to replenish nuclear or cytoplasmic levels resulting in subsequent depletion from both cellular compartments. On the other hand, co-administration of Cdt2 with miR-17∼92 circumvents the need for endogenous mRNA translation. Since miR-17∼92 does not directly target the Cdt2 protein itself, the exogenously administered Cdt2 accumulates in both nuclear and cytoplasmic compartments compensating the miR-17∼92 mediated translational blockage to some extent. Our data indicate that miR-17∼92 treatment results in a significant increase in the S-phase cell population, accompanied by a notable reduction in BrdU incorporation and an increase in apoptotic and sub-G1 cells, highlighting the complex effects of this miRNA cluster on cell proliferation and survival (see Figures 5, 6A–F, S5). These findings suggest that the miR-17∼92 cluster exerts temporal control over the G1/S transition, possibly by maintaining low E2F1 protein levels via MYC regulation, until the G1/S boundary is reached (26,27). Disruption of this regulatory axis leads to premature E2F1 accumulation, activation of a DNA damage-induced checkpoint, and subsequent cell cycle arrest (26–28).

Our observation that miR-17∼92 impedes DNA synthesis, as evidenced by the significant decrease in BrdU incorporation, is further supported by the increase in H2AX phosphorylation (γH2AX) (Figure 6G), a hallmark of DNA damage and replication stress. Notably, the lack of BrdU incorporation in most γH2AX-positive cells suggests that the DNA damage response (DDR) is triggered outside of active DNA synthesis, implicating miR-17∼92 in the induction of replication stress and checkpoint failure (29). This aligns with the emerging view that miRNAs not only fine-tune the timing of cell cycle regulator accumulation but also serve as sentinels for genomic integrity, activating DDR pathways when aberrant cell cycle events occur (27,29). In GBC cells, miR-17∼92-induced destabilization of Cdt2, a key cell cycle regulator, appears to be a major mechanism underlying growth inhibition and impaired metastatic potential. However, the partial rescue of suppressed growth upon Cdt2 co-transfection suggests that miR-17∼92 likely targets additional pathways or factors beyond Cdt2, contributing to its broad anti-tumor activity (28,30). The observed increase in replication stress and DDR markers upon miR-17∼92 treatment further supports the notion that miRNAs act as critical regulators of the cellular alarm system, alerting the cell to oncogenic perturbations and enforcing checkpoints to safeguard genomic stability (27,29). Collectively, our findings position the miR-17∼92 cluster as a potent modulator of cell cycle dynamics, DNA damage response, and cell fate in GBC. The ability of miR-17∼92 to induce apoptosis, inhibit DNA synthesis, and promote replication stress underscores its therapeutic potential as a multifaceted anti-cancer agent. Future studies will be required to delineate the full spectrum of miR-17∼92 targets and to explore the translational relevance of these findings in clinical settings.

Collectively, this study provides comprehensive insights into the genome-wide miRNA landscape for gallbladder cancer, identifying numerous novels and differentially expressed microRNAs in GBC cells. It reveals a complex network of microRNA dysregulation in GBC, with several previously uncharacterized microRNAs exhibiting significant alterations in expression. These newly discovered microRNAs may serve as potential biomarkers for early detection and prognosis of GBC. Upon further study, they may also, uncover functional mechanism underlying their regulatory targets and could give important clues behind carcinogenesis of GBC cells. Moreover, our findings offer a valuable resource for ongoing investigations, with significant potential for enhancing our understanding of disease mechanisms and developing novel diagnostic and therapeutic strategies. Additionally, this study revealed the tumor suppressive role of miR-17∼92 in GBC progression, which acts by directly targeting an oncogenic factor, Cdt2. Investigating the potential of miR-17∼92 *in-vivo* and in clinical settings may ultimately lead to improved patient outcomes and help to develop therapy to curb this deadly disease.

## Declarations Acknowledgement

The authors are thankful to the Director and Dean, Institute of Science, BHU and Coordinator, School of Biotechnology, Institute of Science, Banaras Hindu University for providing space and facilities to conduct the research. We are thankful to the Central Discovery Centre (CDC) and SATHI, BHU for facilitating various facilities. We are also thankful to ISLS, BHU for facilitating the qRT-PCR facility. We would like to extend our gratitude towards Department of Biotechnology (DBT), Govt. of India for funding Samarendra K. Singh (SKS), and Simran Mathur (SM), Council of Scientific and Industrial Research (CSIR), Govt. of India for funding Sonika Kumari Sharma (SS) and University Grant Commission (UGC), Govt. of India for funding Garima Singh (GS). We also thank Department of Science & Technology Fund for Improvement of S&T Infrastructure (DST-FIST), Govt. of India for providing the infrastructure fund to the School of Biotechnology, BHU.

## Authors’ Contribution

Sonika K. Sharma: Investigation, methodology, data analysis, writing original draft, review & editing of the manuscript. Dr. Garima Singh: Helped Sonika K. Sharma in the experiments, data analysis and was involved in critical reviewing and editing of the manuscript. Simran Mathur and Deepanshu Aul: Helped in execution of experiments, data analysis and editing of manuscript. Dr. Vinay K. Singh: Data curation, analysis and uploaded the raw data to database. Dr. Puneet and Dr. Satyendra K. Tiwary: Provided the patient tissue samples. Dr. Samarendra K. Singh: Idea conceptualization, project administration and designing, supervision, critical review, writing & editing of the manuscript.

## Data Availability

The datasets presented and analysed in this study is submitted to the NCBI database. The accession number(s) can be found at: https://www.ncbi.nlm.nih.gov/ with BioProject ID PRJNA1044934. All the data generated and analysed in this study are available from corresponding author on request.

## Ethical Approval

The study was approved by Centre for Genetic Disorder (Ethical board, T.Sc./ECM-XVIII/2024-2025; dated 13/04/2024), Institute of Science, Banaras Hindu University, Varanasi.

## Consent for Participation and Publication

Signed informed consent was obtained from all the patients.

