## Supplementary Figures for "Mapping the miRNA Landscape of Gallbladder Cancer: Genome-Wide Insights and Functional Implications"

**Supplementary Figure 1**

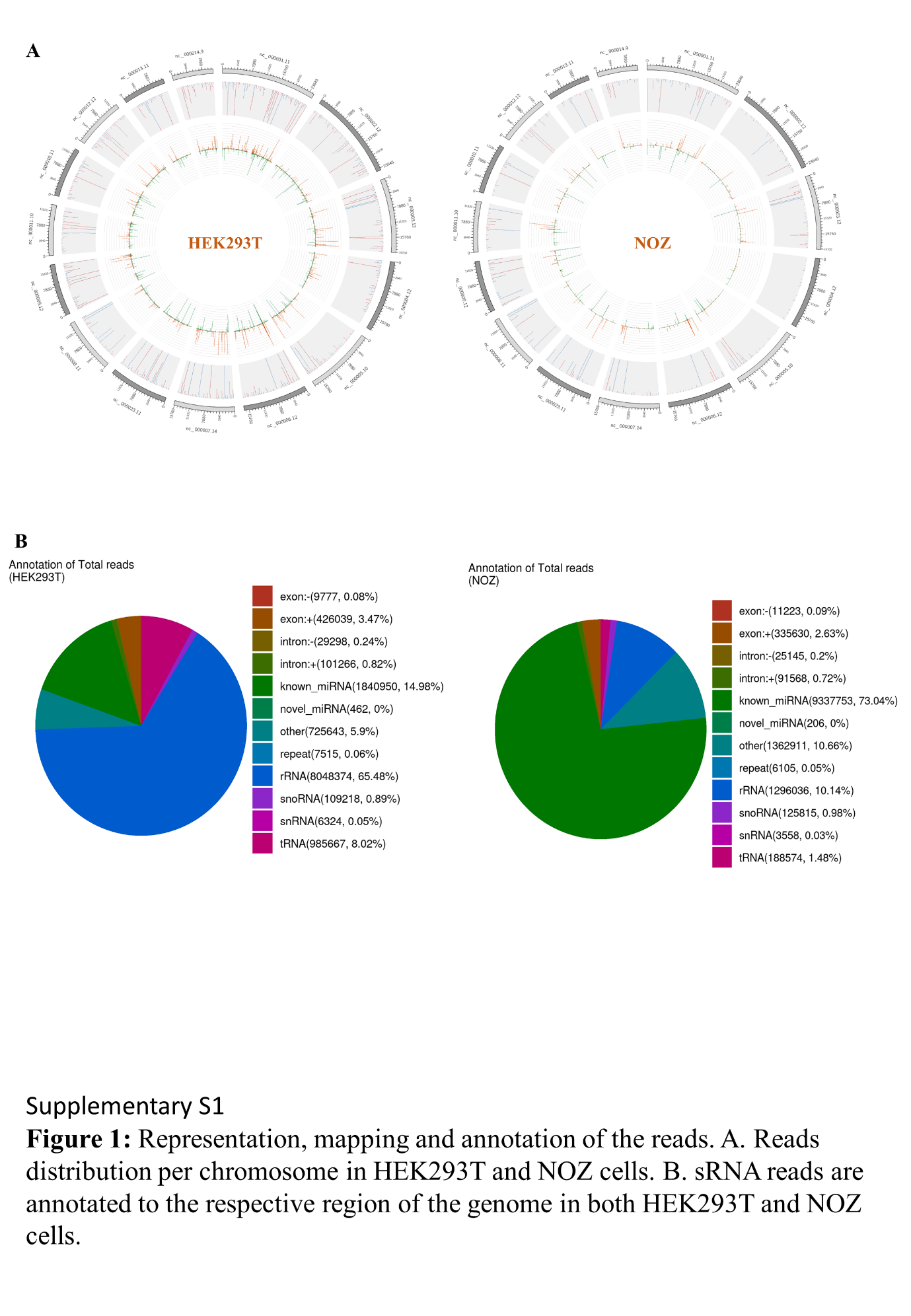

**Figure S1:** Related to Fig. 1A: Representation and annotation of sRNA reads to the respective region of the genome in both HEK293T and NOZ cells.

**Supplementary Figure 2**

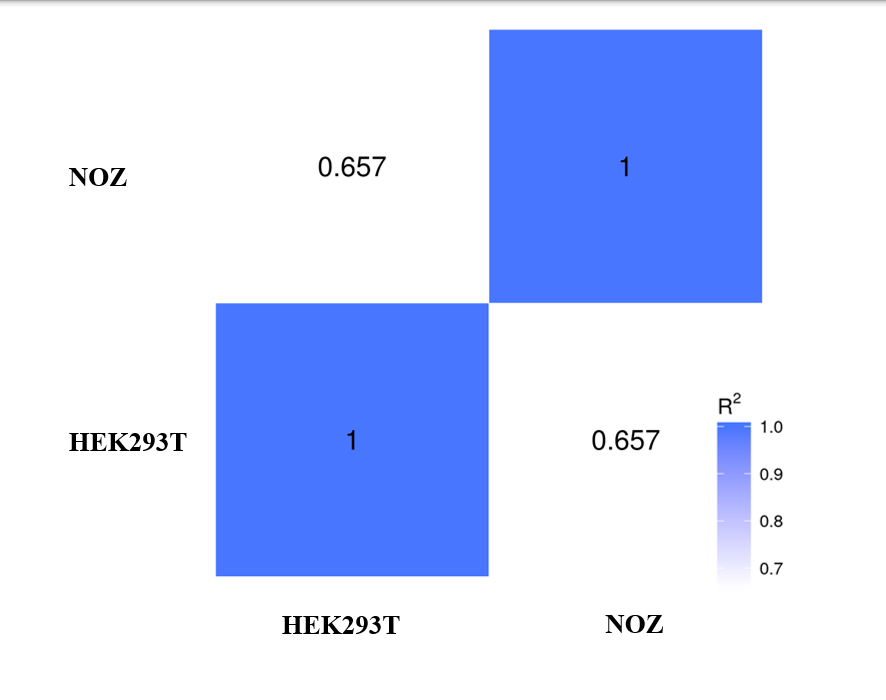

**Figure S2:** Related to Fig. 1C: Pearson correlation between HEK293T and NOZ cells revealing that the two samples are weakly similar to each other with correlation coefficient (R^2^) of 0.657.

**Supplementary Figure 3**

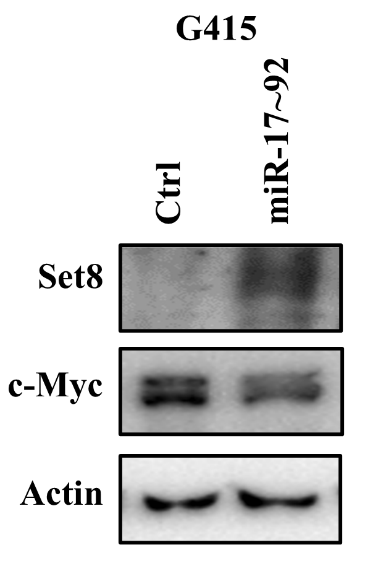

**Figure S3:** Related to Fig. 4F: Western blot analysis for Set8 and c-Myc proteins results in increased level of Set8 and suppressed level of c-Myc in miR-17~92 treated G415 cells.

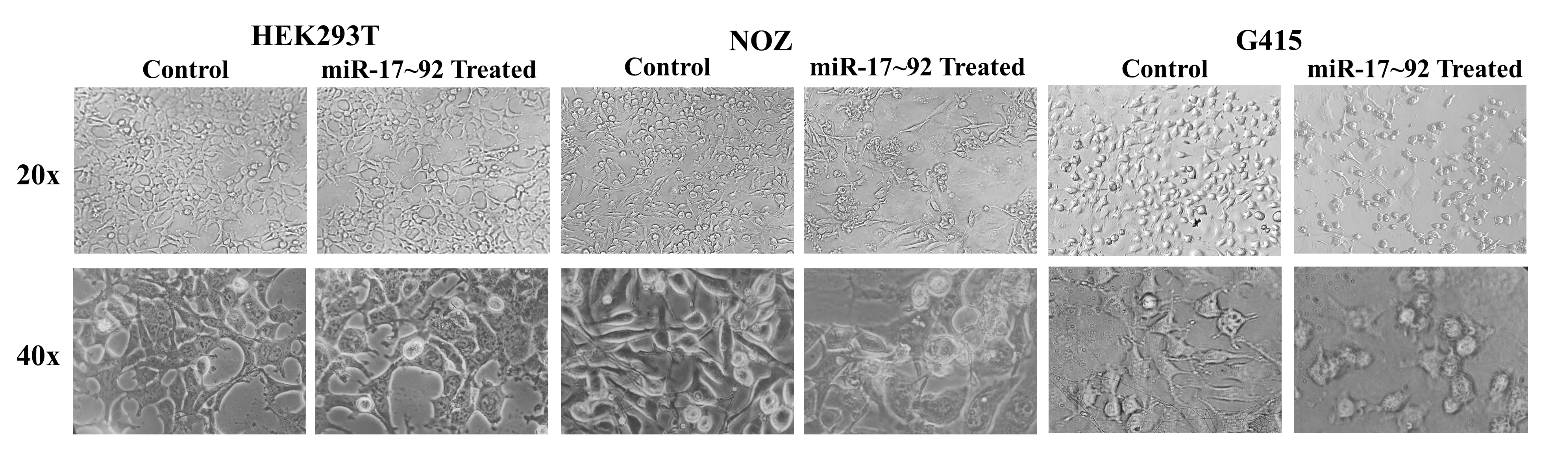
**Supplementary Figure 4**

**Figure S4:** Related to Fig. 5A-C: Phase-contrast microscopic images of HEK293T, NOZ and G415 cells at 4^th^ day after transfection with miR-17~92 at 20x an 40x.

**Supplementary Figure 5**

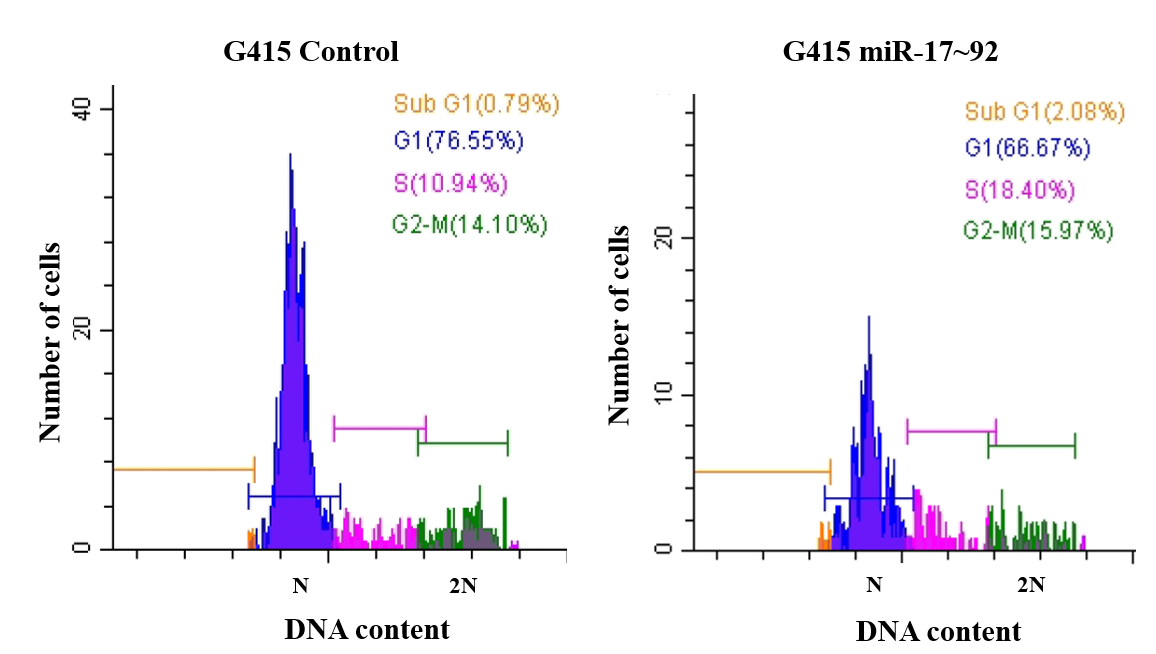

**Figure S5:** Related to Fig. 6A and B: Flow cytometric analysis of Cell cycle representing miR-17~92 induces S-phase arrest in G415 cells.

**Supplementary Figure 6**

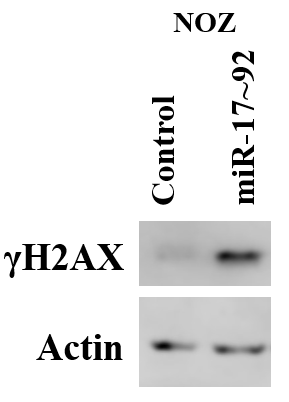

**Figure S6:** Related to Fig. 6E and F: Western blot analysis for γH2AX proteins results in increased level of γH2AX in miR-17~92 treated NOZ cells.

**Supplementary Tables**

**Table 1:** Related to Figure 1: Number of known mature miRNAs and hairpin (pre-miRNA) in each sample.

| **Types** | **Total** | **HEK293T** | **NOZ** |
| --- | --- | --- | --- |
| **Mapped mature** | 1220 | 946 | 971 |
| **Mapped hairpin** | 1065 | 840 | 873 |

**Table 2:** Related to Figure 1: Number of novel mature miRNAs and hairpin (pre-miRNA) in each sample.

| **Types** | **Total** | **HEK293T** | **NOZ** |
| --- | --- | --- | --- |
| **Mapped mature** | 37 | 29 | 27 |
| **Mapped hairpin** | 38 | 31 | 28 |
